# Capillary-scale Microvessel Imaging with High-frequency Ultrasound Localization Microscopy in Mouse Brain

**DOI:** 10.1101/2024.09.19.613950

**Authors:** Matthew R. Lowerison, Yike Wang, Bing-Ze Lin, Zhe Huang, Dongliang Yan, YiRang Shin, Pengfei Song

**Author notes:** Corresponding Author: Dr. Pengfei Song.

## Abstract

Ultrasound localization microscopy is a super-resolution vascular imaging technique which has garnered substantial interest as a tool for small animal neuroimaging, neuroscience research, and the characterization of vascular pathologies. In the pursuit of increasingly high-fidelity reconstructions of microvasculature, there remains several outstanding questions concerning this sub-diffraction imaging technology, including the accurate reconstruction of microvessels approaching the capillary scale and the pragmatic challenges associated with long data acquisition times. In the context of small animal neurovascular imaging, we posit that increasing the ultrasound imaging frequency is a straightforward approach to enable higher concentrations of microbubble contrast agents, thus increasing the likelihood of microvascular/capillary mapping and decreasing the imaging duration. We demonstrate that higher frequency imaging results in improved ULM fidelity and more efficient microbubble localization due to a smaller microbubble point-spread function that is easier to localize, and which can achieve a higher localizable concentration within the same unit volume of tissue. A select example of *in vivo* capillary-level vascular reconstruction is demonstrated for the highest frequency imaging probe, which has substantial implications for neuroscientists investigating microvascular function in disease states, regulation, and brain development. High frequency ULM yielding a spatial resolution of 7.1μm, as measured by Fourier ring correlation, throughout the entire depth of the brain, highlighting this technology as a highly relevant tool for neuroimaging research.

## 1. Introduction

Ultrasound localization microscopy (ULM) is a super-resolution imaging technology which has rapidly gained traction for small animal neurovascular imaging (Couture et al., 2018; Song et al., 2023). First proposed more than a decade ago (Couture et al., 2011; Siepmann et al., 2011), ULM exploits intravascular microbubble (MB) contrast agents to reconstruct vasculature at an imaging resolution below the diffraction limit (Christensen-Jeffries et al., 2015; Errico et al., 2015). The refinement and optimization of ULM processing is an ongoing and rapidly developing area of research; but generally, ULM relies on the sub-wavelength localization of isolated MBs. This necessitates spatially sparse MBs in an imaging frame to avoid interference from overlapping or adjacent MBs. These MB localizations are then tracked frame-to-frame to generate MB trajectories which provide super-resolution vascularity maps as well as physiological indices of blood flow dynamics, including velocity (Christensen-Jeffries et al., 2015; Errico et al., 2015), tortuosity (Shelton et al., 2015), and pulsatility (Bourquin et al., 2022). A consequence of the strategy of relying on sparse MB concentrations is that the technology requires long imaging durations to gradually populate a vascular map (Christensen-Jeffries et al., 2019; Hingot et al., 2019; Lowerison et al., 2020). This in turn is a substantial barrier to the technique, as it necessitates expensive data acquisition practices and requires minimal tissue motion.

Despite these pragmatic concerns, numerous research groups have been actively developing and applying ULM to neuroimaging and to a wide range of neurovascular pathologies. There has been a plethora of demonstrations of the technology in imaging neuroscience including investigation into neurovascular coupling (Renaudin et al., 2022), the effect of aging on neurovasculature (Lowerison et al., 2022), neurodegenerative diseases such as Alzheimer’s disease (Lin et al., 2024; Lowerison et al., 2024), hydrocephaly (Zhang et al., 2022), and stroke in rodent models (Chavignon et al., 2022) and in clinic (Demené et al., 2021). Ultimately, the end goal of ULM research is to provide meaningful physiological indices of microvascular function that are either not possible or not pragmatic with other imaging technologies.

The preeminent accomplishment of a microvascular neuroimaging technology would be the successful mapping of the smallest vessels present (i.e., capillaries) with a high degree of confidence and fidelity throughout the entire depth of the brain. ULM side-steps the conventional trade-off between resolution and penetration depth in ultrasound: the localization process substantially improves resolution without sacrificing the imaging field-of-view (Couture et al., 2018). Under the assumption of perfect imaging conditions (e.g., no tissue motion, no aberration, no MB signal overlap/interference, sufficient signal-to-noise ratio), the ideal ULM localization accuracy is determined by the Cramér-Rao lower bound (Desailly et al., 2015), which predicts a sub-capillary resolution even at clinical imaging frequencies. Empirically, a more realistic localization accuracy *in vivo* tends to be on the order of one tenth of the wavelength (Desailly et al., 2013; Hingot et al., 2021), arising from non-ideal imaging conditions such as high tissue motion and imperfect speed-of-sound estimation. This empirical limit should be sufficiently precise to resolve capillary-scale microvessels for small animal imaging, however the stochastic nature of ULM makes MB traversal for a specific capillary incredibly rare (Christensen-Jeffries et al., 2019; Hingot et al., 2019; Lowerison et al., 2020), reducing the probability and confidence for capillary imaging with ULM. The combination of the pursuit of small vessel mapping and the stochastic nature of ULM also exacerbates the requirement for prohibitively long data acquisition times.

A potential remedy to rare MB traversal events in small vessels and capillaries is to increase the concentration of MBs, resulting in more potential traversal events and therefore a higher probability of reconstructing capillary flow. However, this is at odds with the requirement for spatially sparse MB signals. This problem has motivated several technical developments to extract meaningful MB localization data from overlapping signals in the computational/post-processing domain. These include Fourier-domain filters to synthetically separate MBs (Huang et al., 2020), compressed sensing (Kim et al., 2021), sparse image recovery (Bar-Zion et al., 2018; Yan et al., 2022), and several demonstrations of deep-learning to extract MB locations (Chen et al., 2022; Shin et al., 2024; van Sloun et al., 2021). These techniques offer powerful, and data driven approaches to mitigate the high-density localization problem, which may be combined with strategies for better localization in the data acquisition/imaging domain.

We posit that one such strategy for better MB localization in the imaging domain is increasing the ultrasound imaging frequency. For small animal neuroimaging there is ample room to explore high(er) imaging frequencies (e.g., 30 MHz) due to the relatively shallow imaging target and the strong acoustic backscatter of MB contrast. This provides a straightforward avenue for enabling accurate localization at high(er) concentrations of MBs. MBs are designed to be <5um in diameter to enable vascular passage without risk of embolism. Thus, MBs represent true point scatterers for most physiologically relevant imaging frequencies and should closely resemble the PSF of the ultrasound imaging system. As the imaging frequency increases, the physical extent of the PSF will narrow, and more MBs can occupy the same unit volume while remaining localizable (Belgharbi et al., 2023; Christensen-Jeffries et al., 2019).

To this end, we investigated the use of a high frequency ultrasound imaging transducer (center transmit of 31 MHz) for ULM imaging of mouse brain vasculature. We compared the ULM imaging performance with two lower frequency transducers (15 MHz and 23 MHz center transmit) under conditions of low and high MB concentrations. All other imaging acquisition and experimental parameters were kept as consistent as possible to minimize confounding sources of reconstruction variance. We demonstrate that a higher imaging frequency yields finer vascular reconstruction fidelity, especially in the thalamic and hippocampal regions. We also demonstrate that the highest frequency probe enabled higher MB concentrations and evidence of capillary-level reconstruction. The high frequency imaging was further tested on a Vantage NXT system which provides a higher sampling rate that alleviates the issue of interleaved sampling on the Vantage 256 system. This provides a higher attainable imaging framerate which is beneficial for the ULM tracking step. This dataset was able to achieve a resolution of 7.1μm, as measured by Fourier ring correlation, for the entire depth of the brain. This has substantial implications for neuroscience, as microvascular/capillary imaging is a crucial tool for understanding neurodegenerative diseases, brain development, and neuro-regulation.

## 2. Methods

### 2.1. Ethics statement

All procedures conducted on mice presented in this article were approved by the Institutional Animal Care and Use Committee (IACUC) at the University of Illinois Urbana-Champaign (protocol #22033). All presented experiments were conducted in accordance with the IACUC guidelines. Mice were housed in an animal care facility approved by the Association for Assessment and Accreditation of Laboratory Animal Care. Every attempt was made to minimize the number of animals used and to reduce suffering at all stages of the study.

### 2.2. Animal model

Mouse anesthesia was initialized using a gas induction chamber supplying 4% isoflurane mixed with oxygen. Anesthesia was maintained via a nose cone supplying 2% isoflurane for all procedures. Once anesthetized, the mouse was transferred to a stereotaxic imaging stage (Model 900-U, David Kopf Instruments, Tujunga, CA, USA), and its head was secured in place via ear bars. A cranial window spanning from bregma to lambda was opened in the skull using a rotary tool (Model K1070, Foredom, Bethel, CT, USA) to expose the bilateral expanse of the cerebral cortex. Animal body temperature was maintained at 37°C using a temperature controller (TCAT-2, Physitemp Instruments, Clifton, NJ, USA). A schematic of this imaging setup is demonstrated in **Figure 1**.

**Figure 1.**
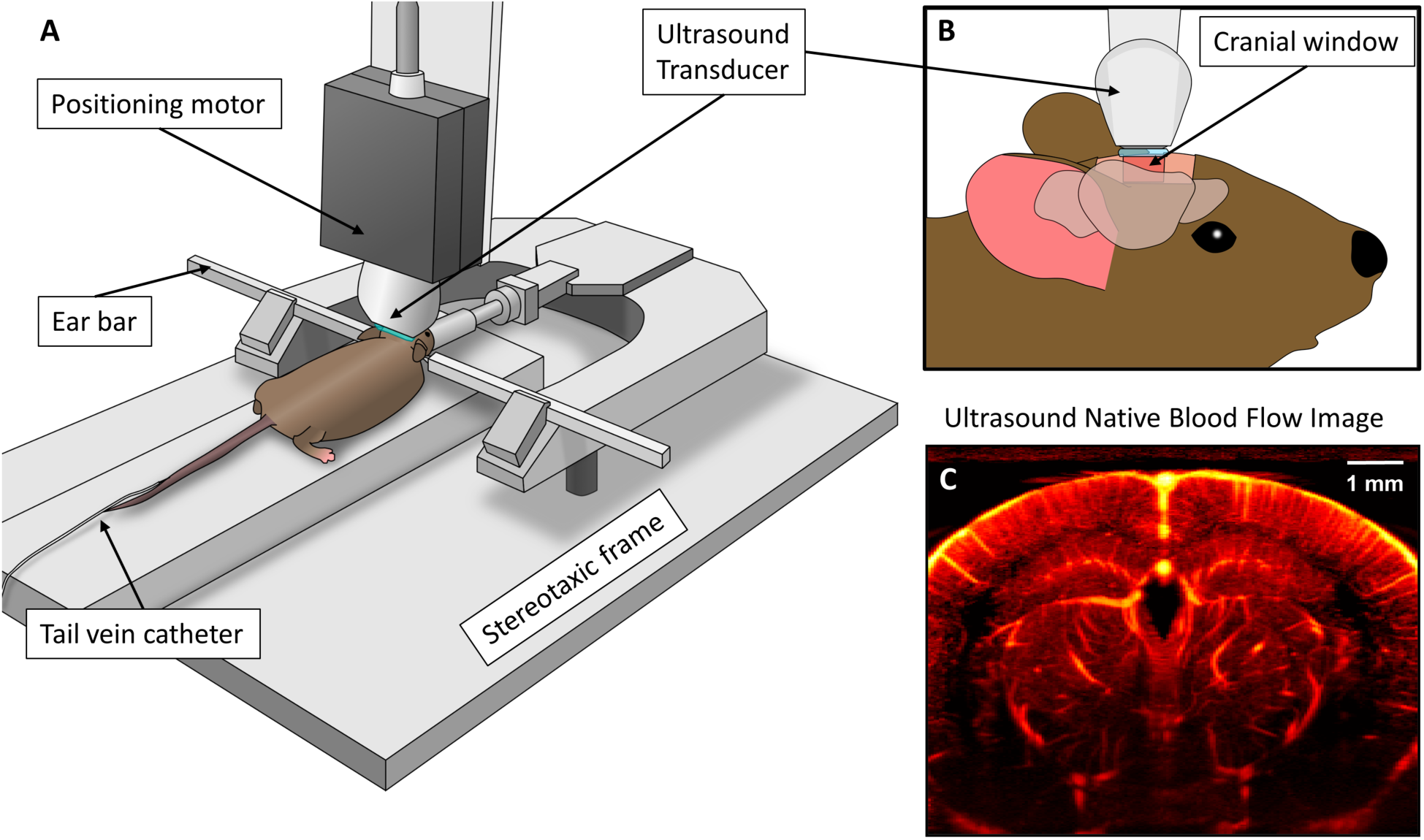
**A)** Schematic of experimental setup, including stereotaxic imaging frame and positioning motor for placement of ultrasound transducer. **B)** Side view of ultrasound imaging through rectangular cranial window opened in the mouse skull, with the imaging plane positioned at approximately 2mm caudal to bregma. **C)** Example ultrasound power Doppler blood flow image, demonstrating select anatomies of interest, such as the hippocampus, thalamus, and cortical regions.

A 29-gauge needle-tip catheter (CA-0099EO, Braintree Scientific, MA, USA) was inserted into the tail vein of the mouse and vessel patency was confirmed with a 50μL saline bolus. Commercially available MB contrast agent (Lumason^®^, Bracco, Milan, Italy) was activated following the supplier’s direction, yielding a solution with approximately 3.5 x 10^8^ MBs/mL. This MB solution was infused through the tail vein catheter at a rate of 20μL/min using a programmable syringe pump (NE-300, New Era Pump Systems Inc., Farmingdale, NY, USA). The MB solution was mixed every three minutes using a custom-built magnetic stirrer to keep the MB concentration consistent throughout the experiment. High MB concentration data acquisitions were accomplished by increasing the flow rate to 40μL/min.

### 2.3. Ultrasound data acquisition

The majority of ultrasound data collection was performed using a Vantage 256 system (Verasonics Inc., Kirkland, WA, USA), with an additional high frequency dataset acquired using a Vantage 256 NXT system (Verasonics Inc., Kirkland, WA, USA). Three different linear array ultrasound transducers were used for imaging: an L22-14vX (Verasonics Inc., Kirkland, WA, USA), an L35-16vX (Verasonics Inc., Kirkland, WA, USA), and an MS-550S (FUJIFILM VisualSonics, Toronto, Canada). Details about these arrays can be found in the following table (Table 1), where the elevational beam width was estimated using the methods described in Section 2.6. All imaging was performed with 9-angle plane-wave compounding (-4 to 4, 1-degree increment), a 1000 Hz post-compounding frame rate, and a one-cycle transmit pulse. A 2-1 interleaved sampling strategy (‘NS200BWI’) was employed to effectively double the Vantage’s 62.5 MHz ADC receive sampling rate for the L35-16vX and MS-550S acquisitions. High frequency imaging was also performed with a Vantage NXT system (Verasonics Inc., Kirkland, WA, USA), where no interleaved sampling was necessary because of the higher system sampling frequency (125 MHz).

**Table 1.**
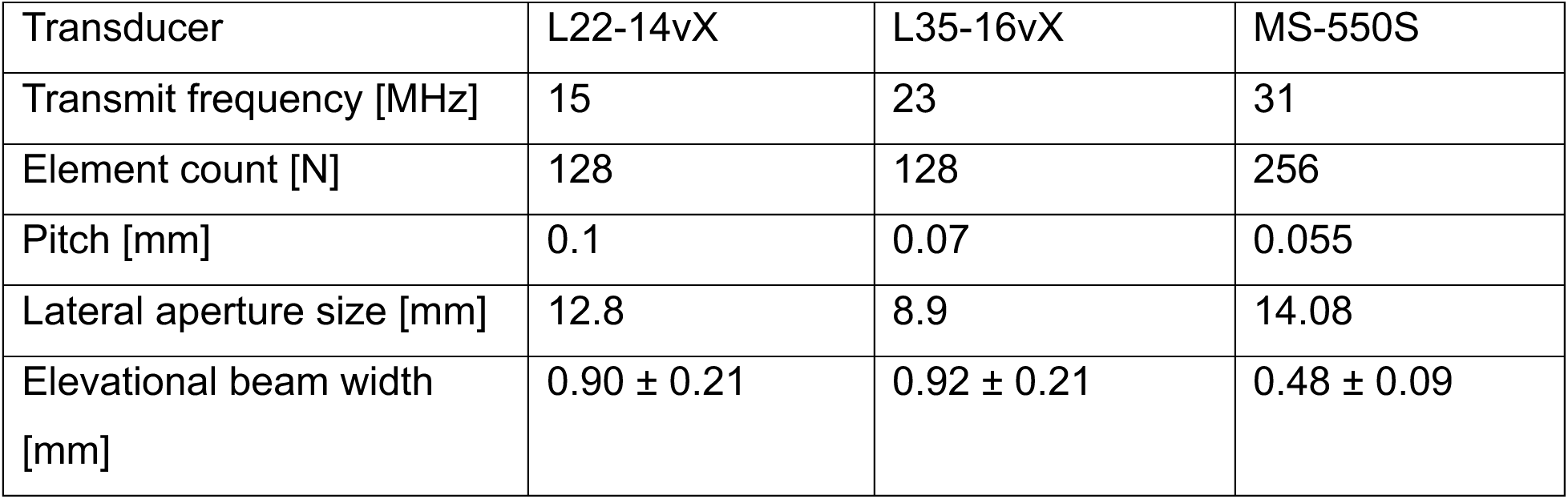
Transducer specifications for L22-14vX, L35-16vX, and MS550-S.

| Transducer | L22-14vX | L35-16vX | MS-550S |
| --- | --- | --- | --- |
| Transmit frequency [MHz] | 15 | 23 | 31 |
| Element count [N] | 128 | 128 | 256 |
| Pitch [mm] | 0.1 | 0.07 | 0.055 |
| Lateral aperture size [mm] | 12.8 | 8.9 | 14.08 |
| Elevational beam width [mm] | 0.90 ± 0.21 | 0.92 ± 0.21 | 0.48 ± 0.09 |

Each ultrasound transducer was secured to a translational motor (VT-80 linear stage, Physik Instrumente, Auburn, MA) that was part of the stereotaxic imaging frame via a 3D-printed transducer holder (**Figure 1A**). Ultrasound gel was applied directly to the surface of the brain, and the transducer was coupled to produce a coronal anatomical imaging plane. The motorized stage was then adjusted to find an imaging position that was at approximately bregma -2 mm. Care was taken to attempt to align each of the different transducers with the same anatomical imaging plane, using a combination of translational motor positions and qualitative assessment via ultrasound blood flow imaging (**Figure 1C**).

For each dataset, a total acquisition time of 300 seconds (300,000 frames) of contrast-enhanced ultrasound data was acquired. Ultrasound data was stored as raw radiofrequency (RF) datasets for offline beamforming. Beamformed images were saved as in-phase/quadrature (IQ) data for further processing.

### 2.4. ULM reconstruction

All ULM reconstruction and analysis was performed in MATLAB (The MathWorks, Natick, MA; version R2022b). MB signal was extracted from the contrast-enhanced IQ data using singular value decomposition-based clutter filtering (Desailly et al., 2017), where the low-order tissue threshold was determined adaptively (Song et al., 2017b). This generally selected the first 20 singular values to be excluded. A noise-equalization profile was then applied (Song et al., 2017a), and a MB separation filter (Huang et al., 2020) was used to split the MB data into upward and downward components.

The filtered IQ data was then spatially interpolated to an isotropic λ/10 grid (L22-14vX: 9.8μm; L35-16vX: 6.6μm; MS-550S: 4.9μm) and normalized cross-correlation was performed with an empirically determined MB template (2D Gaussian) to produce sub-pixel localization candidates. These candidates were then paired and tracked using the uTrack algorithm (Jaqaman et al., 2008) with a minimum persistence of 10 frames (10 ms). MB trajectories were then plotted onto a reconstruction grid space with a pixel size 2μm and accumulated into a final super-resolved vascular image.

### 2.5. ULM image analysis

ULM image resolution was estimated using Fourier Ring Correlation (FRC) following the protocol outlined by (Hingot et al., 2021). Briefly, each MB track was randomly assigned to one of two ULM sub-accumulations, reconstructed with a pixel size of 1μm, to produce independent reconstructions of the same vasculature. These two images were Fourier transformed and the correlation between the two spectra was calculated for rings of increasing frequency to produce an FRC curve. The resolution of image was estimated using the two standards typically applied to ULM characterization: the ½-bit threshold and 2-σ criterion (van Heel & Schatz, 2005). The saturation rate of ULM images was quantified using an exponential saturation model as described by (Dencks et al., 2017; Lowerison et al., 2020):

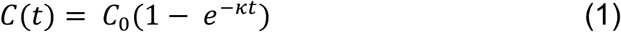

where *C*_0_is the max saturation level of the curve (assumed to be 100% of the vasculature), t is time in seconds, and κ is a rate constant. The characteristic time was taken as the estimate to reach a 90% mapping of microvessels.

### 2.6. Transducer elevational beam width estimation

The elevational beam width of each transducer was estimated by scanning a zero-degree planewave transmit with a capsule hydrophone (HGL-0200, Onda Corporation, Sunnyvale, CA, USA) and ONDA AIMS III system (Onda Corporation) in a water tank. The ONDA position system was used to move the hydrophone in the axial direction starting from 2mm away from the transducer surface to a maximum depth of 8mm (0.05mm increment) and from -2mm to 2mm in the elevational direction (0.02 mm increment). Elevational beamwidth was estimated as the full width at half max of the root mean square voltage at each axial position.

## 3. Results

### 3.1. MB PSF has reduced physical extent at higher frequency

With the Allen Brain Atlas to provide anatomical context (**Figure 2A**), example MB contrast-enhanced imaging frames for the three different transducers are demonstrated in **Figure 2B**, along with zoomed-in insets to identify individual isolated MBs. Three anatomical regions of interest are highlighted, corresponding to the cortex (white box), hippocampus (green box), and thalamus (red box). It is worth noting that the local concentration of MBs varies substantially between these different regions, with the thalamus having a much higher concentration than the cortex. As the imaging frequency is increased, the spatial extent of the MB PSF is reduced, potentially allowing for more efficient localization at higher MB concentrations. This is evident when comparing the thalamic region of the L22-14vX image, which has a substantial amount of MB PSF overlap in comparison to the MS-550S image, where the majority of MBs remain as isolated point targets.

**Figure 2.**
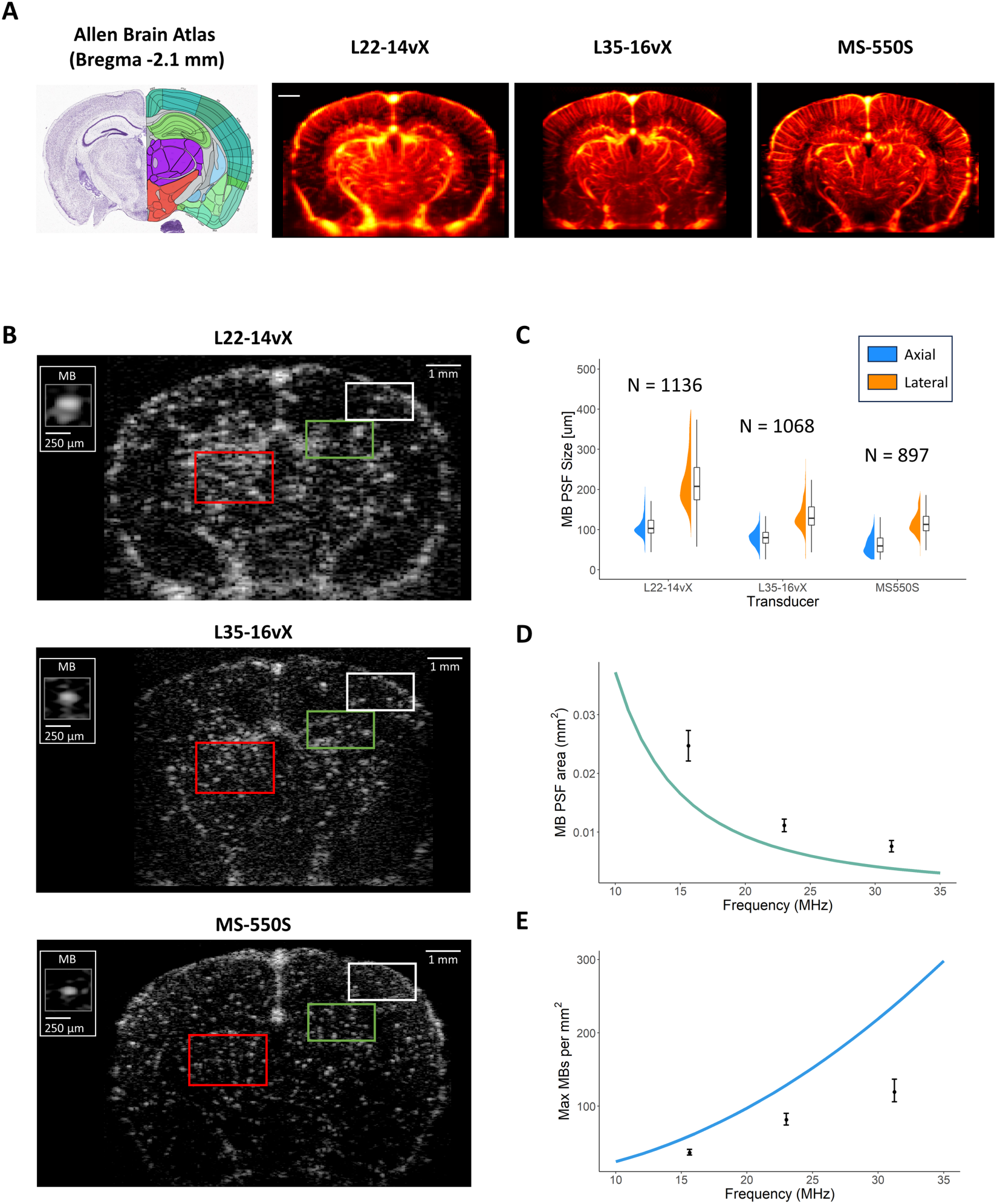
**A)** Reference Allen Brain Atlas coronal section along with example of SVD clutter-filtered contrast-enhanced accumulations for the three different transducers. **B)** The MB PSF has a reduced physical extent as the imaging frequency is increased, as shown by the figure insets demonstrating an isolated MB for each transducer. Three brain anatomies of interest are highlighted in the ultrasound images, the cortex (white box), the hippocampus (green box), and the thalamus (red box), which demonstrate different local concentrations of MBs. There is substantial MB PSF overlap at lower frequency that is resolvable at higher frequency. **C)** Experimental measurements of MB PSF sizes in both the axial and lateral dimensions for the three transducers. **D)** A theoretical plot of MB PSF cross-sectional area with respect to imaging frequency, along with the experimental measurements for each transducer. **E)** A hypothetical plot of the maximum number of resolvable MBs per square millimeter, with experimental estimates.

In **Figure 2C** we demonstrate experimentally determined MB PSF sizes, reported as the full width at half max in both the axial and lateral dimensions. For the L22-14vX, the MB PSFs were 113.0 ± 39.2μm axial by 218.8 ± 66.3μm lateral (number of isolated MBs analyzed N = 1136); for the L35-16vX, they were 80.9 ± 22.8μm axial by 137.7 ± 47.4μm lateral (N = 1068); and for the MS-550S, they were 63.7 ± 26.1μm axial by 119.1 ± 36.8μm lateral (N = 897). A rough approximation for the MB PSF cross-sectional area under these conditions is to consider it as an ellipse with an axial dimension equal to the wavelength and a lateral dimension of two wavelengths. This is demonstrated in the plot in **Figure 2D**, for frequencies ranging from 10 to 35 MHz, under the assumption of an average tissue sound speed of 1540 m/s. The experimentally determined MB PSF areas are included as points in this graph. To get an idea of how MB localization efficiency can improve with increasing frequency, we have plotted a theoretical “maximum” MB concentration in terms of resolvable MBs per millimeter squared of tissue, **Figure 2E**, estimated using the MB PSF area and the optimal packing density of elliptical surfaces (Chang & Wang, 2010).

### 3.2. Low MB concentration side-by-side comparison of ULM reconstruction

An example of mouse brain ULM images with the L22-14vX, L35-16vX, and the MS-550S transducers is demonstrated in **Figure 3**, all taken under low MB concentration conditions that are optimized for the lower frequency transducer. Generally, the fidelity of ULM imaging was improved with increased frequency, providing narrower vessel diameters in the cortex (white box) and clearer examples of microvascular branching points in the hippocampus (green box) and thalamus (red box). The improvement in fidelity is particularly evident in the thalamic region, which was previously noted to be a region with high local concentrations of MBs (**Figure 2B**). However, we found that the ULM vessel saturation decreased with increased imaging frequency in the deepest regions of the brain (e.g.: the entorhinal cortex). This can likely be attributed to a combination of narrower elevational beamwidth (**Table 1**), which reduces the sampling volume, and increased signal attenuation for higher frequencies. The estimated 90% saturation times for the L22-14vX, L35-16vX, and the MS-550S were 181.7, 191.1, and 307.2 seconds, respectively.

**Figure 3.**
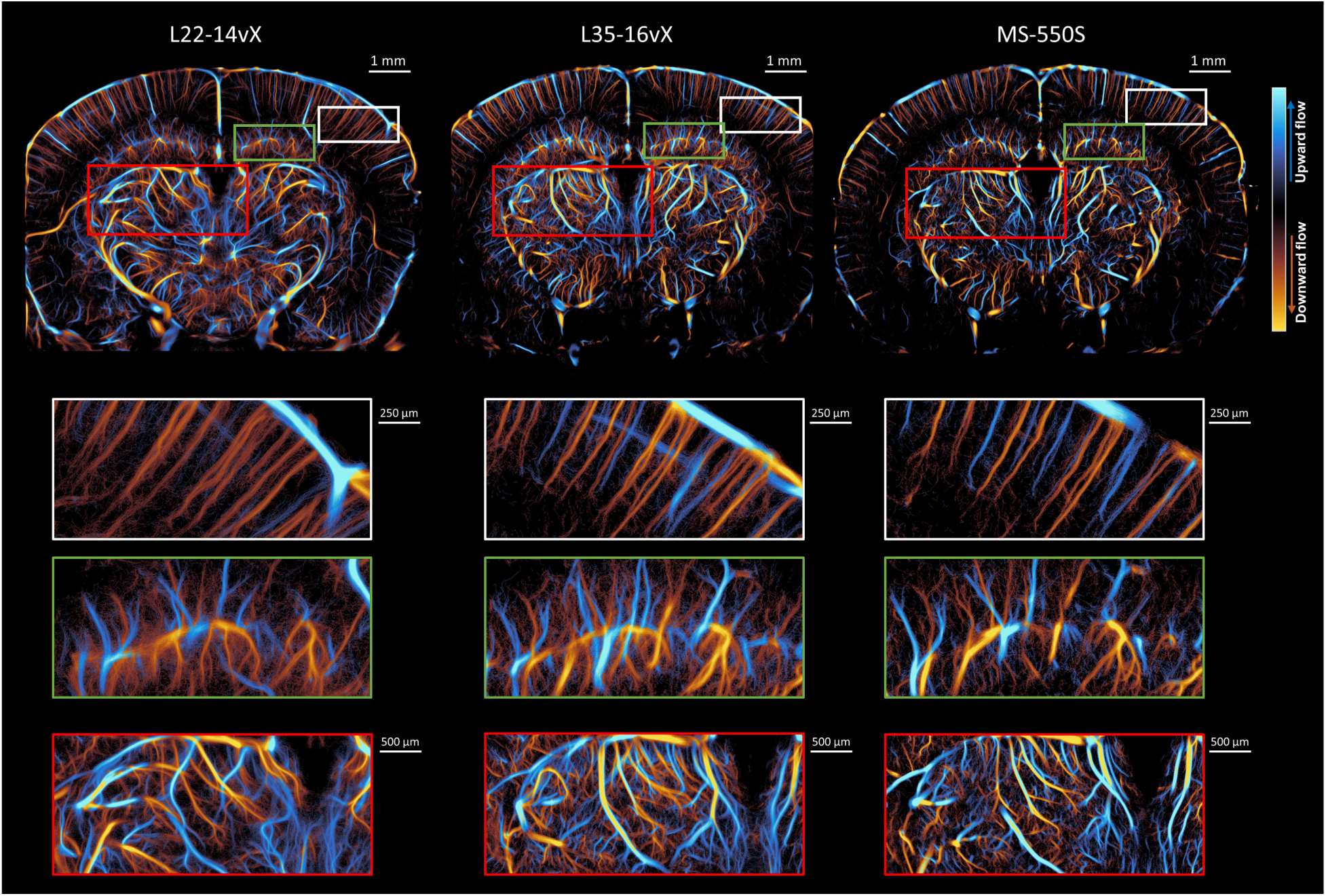
Side-by-side comparisons of the ULM reconstructions for the three different transducers at low MB concentration. The vessel saturation is decreased for increasing frequencies, especially for the deeper regions of the brain such as the entorhinal cortex. The white figure insets highlight the cortical vessel reconstruction, which demonstrated clearer vessel diameters at higher frequency. The green and red insets focus on the hippocampus and thalamus, respectively, where less noisy vascular reconstruction and more evident microvascular branching points are evident at high frequency.

The differences in reconstruction fidelity for this dataset was quantified using Fourier Ring Correlation (**Figure 4**) to estimate the spatial resolution. A diagrammatic example of the FRC process is demonstrated in **Figure 4A**, where individual MB tracks were accumulated into two independent ULM reconstructions. We found that there was a gradual improvement in the ULM resolution with increasing transducer frequency (**Figure 4B-D**). The L22-14vX had a ½-bit resolution of 19.2 μm; the L35-16vX had a ½-bit resolution of 18.1 μm; and the MS-550S had a ½-bit resolution of 14.8 μm. The 2-σ resolutions were 15 μm, 15.8 μm, and 12.3 μm, respectively. It should be noted that the 2-σ threshold is close to the noise floor for ULM imaging, potentially explaining the slight degradation of resolution for the L35-16vX.

**Figure 4.**
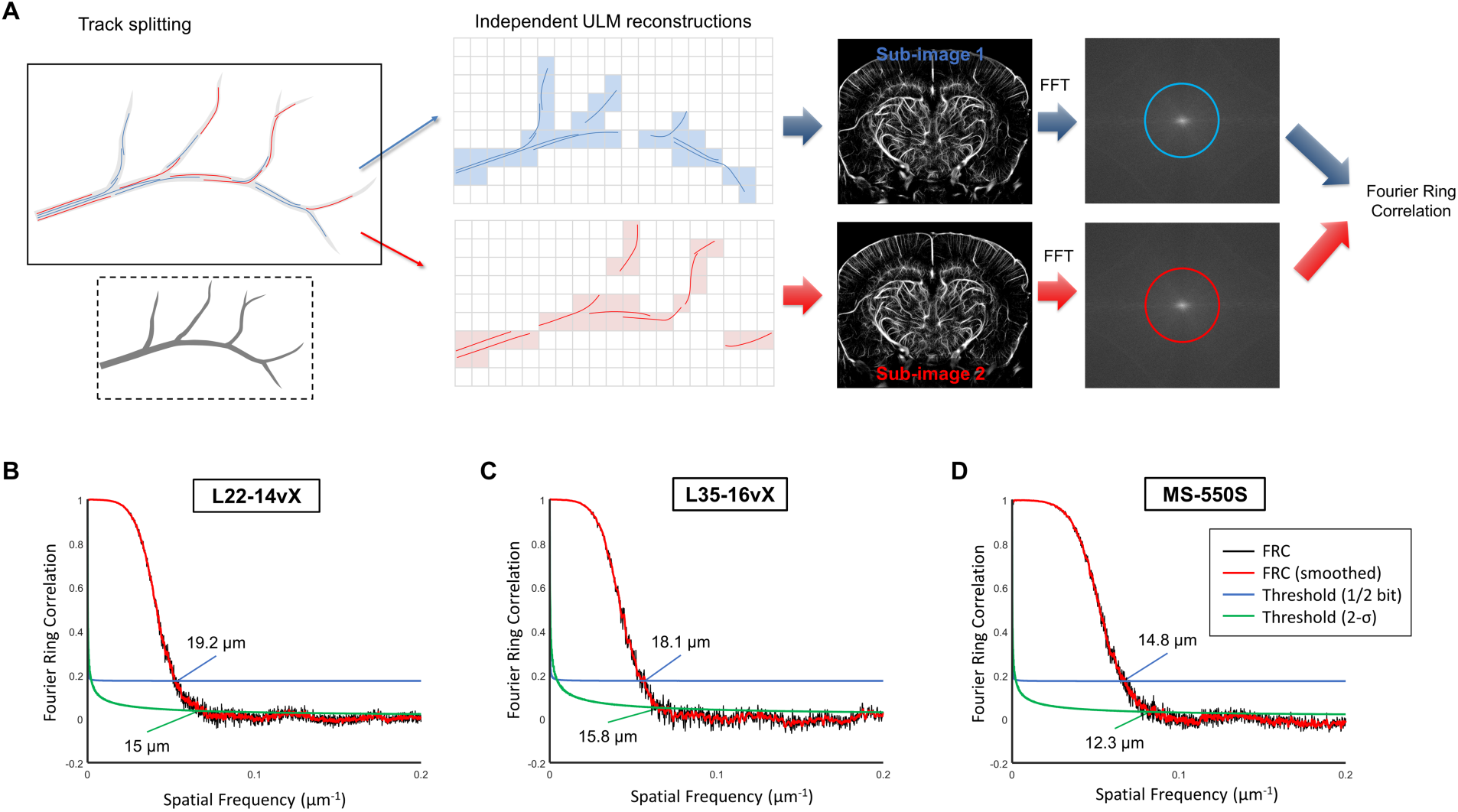
Fourier Ring Correlation resolution estimates for demonstrated low MB concentration datasets. **A)** A diagrammatic example of the FRC process, where MB tracks are split into independent ULM reconstructions and compared. The FRC ½-bit and 2-σ estimates demonstrates a trend of gradually improved ULM resolution with increasing transducer frequency, with the **B**) L22-14vX yielding a resolution estimate of 19.2 / 15 μm, the **C**) L35-16vX an estimate of 18.1 / 15.8 μm, and the **C**) MS-550S an estimate of 14.8 / 12.3 μm.

### 3.3. High MB concentration side-by-side comparison of ULM reconstruction

The same sort of comparison of mouse brain ULM images between the three transducers was also performed at a higher MB concentration (**Figure 5**). The ULM image with the L22-14vX demonstrates a substantial degradation in reconstruction fidelity, which is most pronounced in the thalamic region (red box). The microvascular lattice in this inset is noisy and difficult to visualize, and the larger vessels are overemphasized with indistinct branching points. A similar observation is noted for the L35-16vX in this region, although the degradation is less pronounced. The MS-550S has maintained a high degree of reconstruction fidelity, with better saturation under this higher concentration condition. The performance of the three transducers in the cortical region does not appear to be as impacted, likely due to the lower local concentration of MBs in the cortex noted in **Figure 2**. The estimated 90% saturation times under high MB concentration were 209.8, 175.4, and 169.4 seconds for the L22-14vX, L35-16vX, and the MS-550S, respectively.

**Figure 5.**
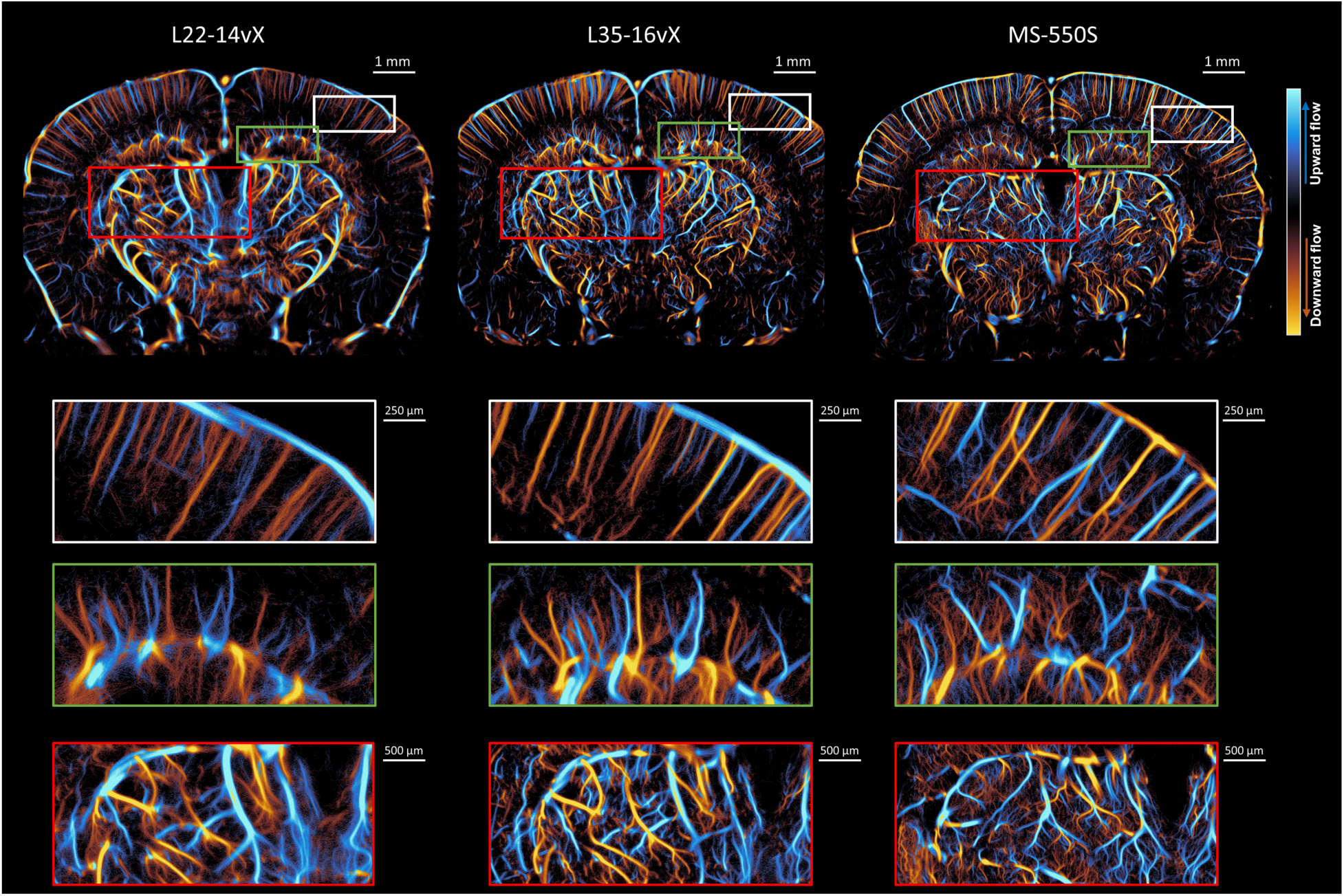
Side-by-side comparisons of the ULM reconstructions for the three different transducers at high MB concentration. The L22-14vX demonstrates a degradation of ULM reconstruction fidelity, especially in the thalamic region (red box), where smaller vessels are difficult to visualize, and larger vessels are overemphasized. A similar observation is noted for the L35-16vX. The MS-550S had better saturation while maintaining a high degree of reconstruction fidelity.

Another mouse was imaged using the MS-550S transducer at high MB concentration using the Verasonics NXT system. This is demonstrated in **Figure 6A**. We noted that the NXT was able to reconstruct more microvasculature in the entorhinal cortex (red box). The hippocampal / white matter region (green box) also demonstrated better saturation with the NXT. This is likely due to a combination of the NXT having a sampling rate that is better suited to imaging with the high frequency MS-550S, and due to the increased acoustic output of the NXT relative to the Vantage 256. The 90% saturation rate for this dataset was estimated at 153.9 seconds.

**Figure 6.**
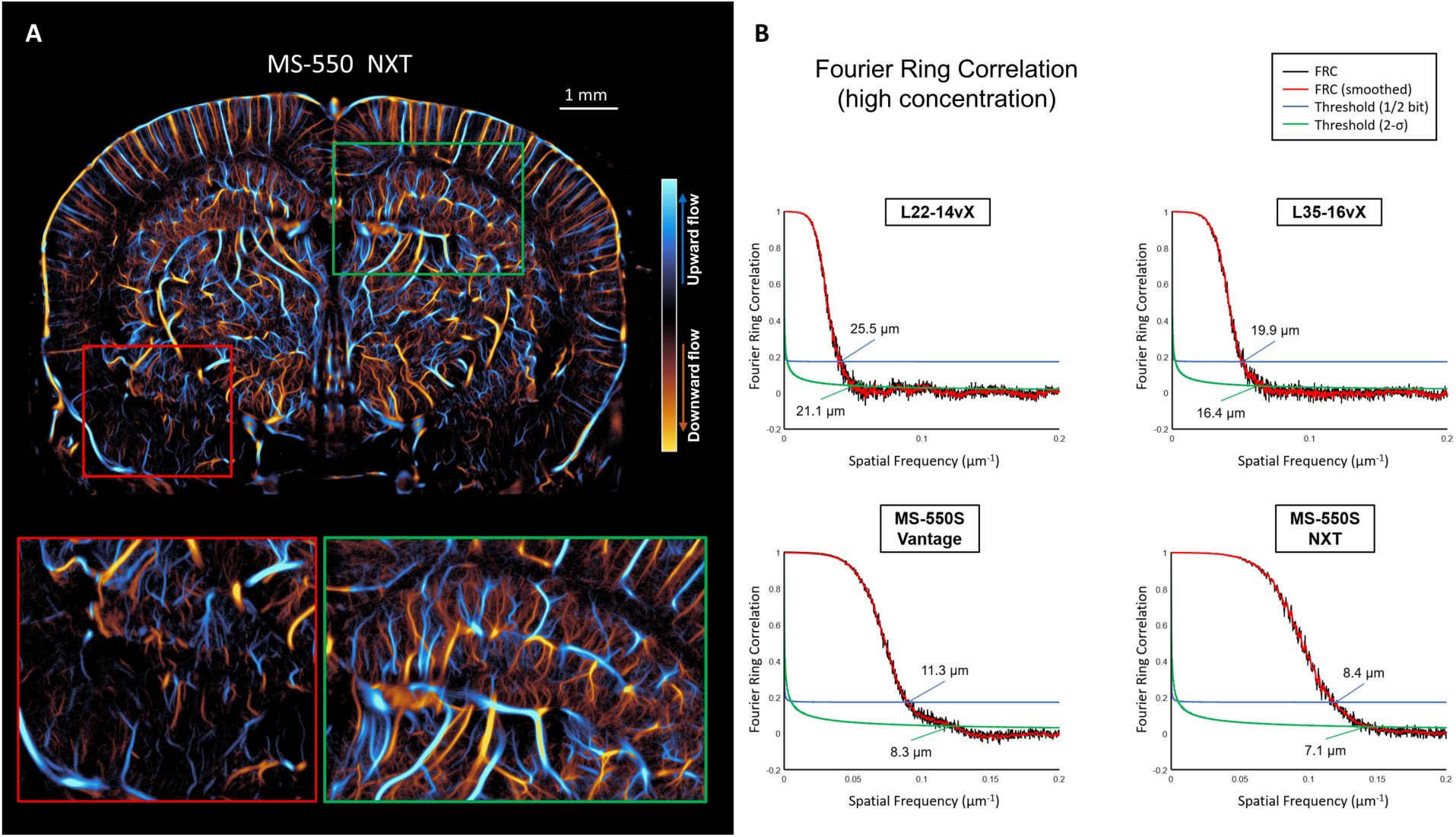
**A)** High frequency imaging was further tested on a Vantage NXT system. This resulted in a high degree of vessel saturation while maintaining reconstruction fidelity. The NXT had better performance for deeper brain regions, such as the entorhinal cortex (red box), than the Vantage 256. The hippocampal / white matter region (green box) also demonstrated better saturation with the NXT. **B)** FRC resolution estimates for the high MB concentration datasets. The L22-14vX had a reduction in the resolution estimate in comparison to the low concentration case, with ½-bit and 2-σ estimates of 25.5 μm and 21.1 μm, respectively. The performance of the L35-16vX was slightly degraded in comparison to the lower concentration case at 19.9 / 16.4 μm. On the Vantage, the MS-550S resolution estimate was improved at higher MB concentration at 11.3 / 8.3 μm. An even finer resolution estimate was achieved with the NXT system, at 8.4 / 7.1 μm.

FRC analysis was also performed on the high MB concentration datasets (**Figure 6B**). The L22-14vX had a reduction in the resolution estimate in comparison to the low concentration case, with ½-bit and 2-σ estimates of 25.5 μm and 21.1 μm, respectively. The performance of the L35-16vX was slightly worse in comparison to the lower concentration case, with a resolution estimate of 19.9 μm (½-bit threshold) and 16.4 μm (2-σ threshold). We found that the Vantage MS-550S FRC analysis yielded a resolution estimate of 11.3 μm (for ½-bit threshold) and 8.3 μm (2-σ threshold). The NXT MS-550S FRC curve estimated a resolution of 8.4 μm (for ½-bit threshold) and 7.1 μm (2-σ threshold). The improvement in ULM resolution relative to low MB concentration datasets can likely be attributed to the increased vessel saturation, providing more instances of reproducible high-spatial frequency information.

### 3.4. Capillary reconstruction via arteriole-to-venule MB transition event

To further investigate the utility of high-frequency ULM imaging, we performed a sequential magnification analysis of a dataset acquired with the MS-550S probe at high MB concentration, which is demonstrated in **Figure 7**. The figure insets display increasing levels of magnification to demonstrate the reconstruction quality for the cortical region. The second magnification inset focuses on a region of the cortex where two counter-directional cortical vessels were extracted to investigate this region in more detail. The main arteriole/venule luminal space is apparent, with some evidence of microvessels branching off to other regions of the cortex. The final magnification demonstrates a “j-shaped” MB trajectory that transitions from the highlighted arteriole to venule in the cortex, implying capillary-level vascular flow. The overlayed arrows denote the direction of MB flow for this trajectory. Other similar trajectory phenotypes are present in the datasets.

**Figure 7.**
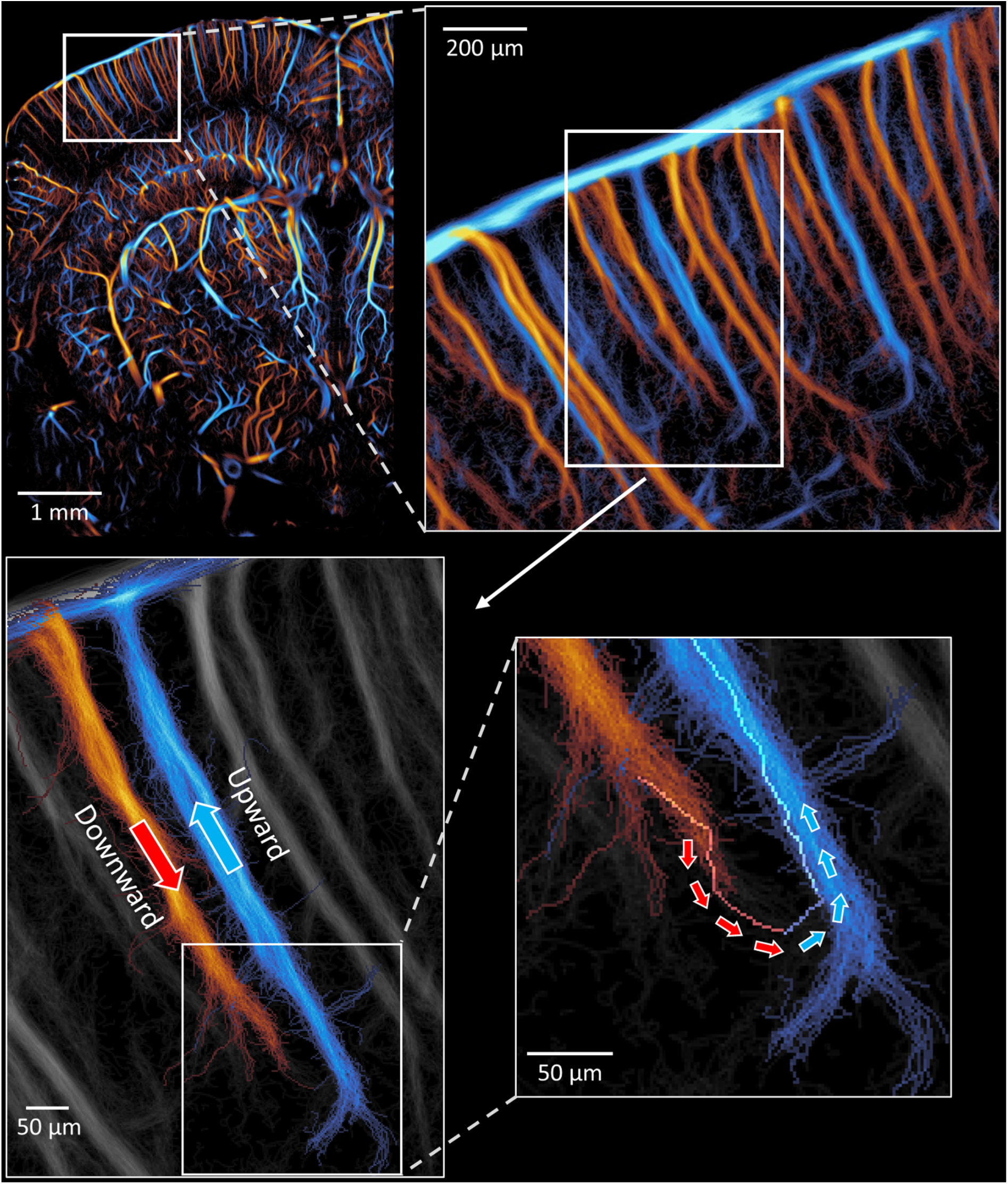
Specific example of arteriole-to-venule transition event. On a separate animal, a high MB concentration dataset was acquired using the MS-550S transducer and the Vantage 256. The figure insets demonstrate a progressive magnification to a cortical region of the brain. Finally, the last magnification level demonstrates a “j-shaped” MB trajectory that exhibited a transition between a cortical arteriole to a cortical venule, implying capillary level vascular reconstruction.

## 4. Discussion / Conclusion

In this report, we investigated the ULM reconstruction fidelity of mouse brain imaging for three ultrasound imaging transducers with different imaging frequencies. We found that increasing the imaging frequency decreased the spatial extent of the MB PSF (**Figure 2**), enabling localization at high(er) MB concentrations. This observation is supported by literature which models the number of MBs that can occupy the same unit volume while remaining localizable (Belgharbi et al., 2023; Christensen-Jeffries et al., 2019). This was particularly relevant for accurate reconstruction of the thalamic regions of the brain, which tended to have higher local concentrations of MBs in comparison to the cortex. One should note that the experimental PSF sizes were larger than the theoretical wavelength for the transducer center frequencies (i.e., 98.6μm at 15.625MHz for the L22-14vX; 66.9μm at 23MHz for the L35-16vX; and 49μm at 31.25MHz for the MS-550S), which can be attributed to factors such as diffraction, low number of compounding angles, and sound-speed inhomogeneities.

For the transducer comparison under the same low MB concentration experimental conditions, we found that the highest frequency probe yielded ULM images with narrower vessel diameters in the cortex, and clearer reconstruction of the microvasculature in the hippocampus and thalamus, in comparison to the lower frequency probes. However, the ULM vessel saturation was reduced for the high frequency probe, which is likely the result of the narrower elevational beamwidth (**Table 1**). This smaller sampling volume can be beneficial for MB localization at higher concentrations: two-dimensional ULM suffers from projection of three-dimensional vasculature (Heiles et al., 2022), leading to overlapping PSFs from distant MBs and elevational ambiguity of MB trajectories. A narrower elevational beamwidth reduces the probability that MBs will overlap, leading to more efficient localization and better elevational resolution. This elevational ambiguity is also problematic for the MB tracking step, as physically distant MBs can be erroneously paired, which can potentially obscure microvascular flow. It is worth noting that previous research has demonstrated that these out-of-plane MB positions can be useful for elevational estimation/visualization in 2D ULM (Renaudin et al., 2023), but this was unexplored in this report.

Under the high(er) MB concentration condition, we found that the ULM reconstruction fidelity of the lowest frequency probe was substantially degraded, especially in the thalamic regions of the brain: the microvascular bed was indistinct / obscure, and the larger vessels lacked obvious branching points. The L35-16vX was less impacted, however the major vessels appear to be overemphasized relative to the low MB concentration reconstruction. Notably, these regions still have a blurry “vessel-like” appearance, with some noisy background haze, which may stem from false localizations of MB interference patterns. These interference patterns will tend to create local maxima that are situated close to the vessel lumen, but which lack the precision and consistency of true MB point localizations. In comparison, the ULM reconstruction fidelity was maintained for the highest frequency probe, with an increase in the overall saturation, especially for the dataset acquired with the NXT system.

The FRC analysis demonstrated that the estimate for the ULM reconstruction resolution modestly improved with increasing frequency under the low MB concentration condition, from 15 μm at the lowest frequency up to 12.3 μm at the highest frequency. For the higher MB concentration datasets, we found that the lowest frequency probe had a worse resolution estimate (21.1 μm) and a marginal decrease in performance for the mid-frequency probe (16.4 μm). For the highest frequency probe, the MS-550S transducer, the FRC resolution estimate improved drastically to 8.3 μm (Vantage 256) and 7.1 μm (NXT). This substantial improvement in the high concentration MS-550S result in comparison to the low concentration MS-550S result is probably attributable to the increase in vessel saturation. FRC is ultimately a measure of consistency between the two independent ULM reconstructions. For the low concentration MS-550S ULM reconstruction, there are likely microvascular MB trajectories which are only present in one of the two sub-images used for FRC analysis. This would lead to these trajectories being discounted as reconstruction noise on the FRC curve. With the higher MB concentration data, there are more overall microvascular trajectories being accumulated, increasing the consistency of the FRC sub-images, and thus improving the quantified spatial resolution. This provides evidence to support our initial hypothesis that increasing the imaging frequency provides an avenue for enabling ULM at high(er) concentrations of MBs, and thus reduces the total accumulation time required for microvascular reconstruction.

As part of this investigation, we demonstrated an exemplary case of an arteriole-to-venule MB transition event in the cortex of a mouse brain, which implies capillary reconstruction fidelity. It should be noted that we are not the first to demonstrate such “j-shaped” MB trajectory phenotypes (Chabouh et al., 2024; Lee et al., 2024; Renaudin et al., 2022), and the limitations of such an example should be discussed. First, this example was manually selected from a list of raw MB tracks, which introduces a substantial risk of investigator bias. The manual selection process is also laborious and under-represents the prevalence of these types of trajectories. This ties into the stochastic nature of ULM imaging: in principle these microvascular trajectories should be in almost all ULM reconstructions, but there is no guarantee that they will present with a compelling phenotype. Second, this “j-shaped” phenotype merely implies capillary/microvascular flow, there is no ground-truth available to validate these findings. The validation of ULM vascular reconstruction via other imaging modalities is an active area of research, and external validation of capillary-level reconstruction remains elusive.

There are other limitations which should also be discussed. For this report we limited our discussion to neuroimaging in small animal models, where the imaging depth is shallow and imaging frequency is usually >15 MHz, thus the empirical limit of one tenth of the wavelength (Desailly et al., 2013; Hingot et al., 2021) should be sufficiently precise to resolve capillary-scale microvessels. It would be challenging to observe these microvascular flow events in larger animals and in humans. High frequency imaging has higher signal attenuation, which limits MB detection in deep regions. Increasing the transmit power can help in this scenario, however there is a risk of more native red blood cell backscatter in the near field (Kuo & Shung, 1994), which can lead to incorrect localizations in the shallow regions of the brain. Higher frequency imaging also requires a higher data sampling rate, which can increase the raw data size and computational requirements. The discussion of the ultrasound PSF with respect to the imaging frequency was simplified for brevity/clarity in the initial hypothesis. Ultrasound has an anisotropic and spatially variant PSF. For planewave compounding the axial imaging resolution depends on the center frequency, bandwidth, and number of cycles of the transmit pulse. The lateral resolution depends on the size of the lateral aperture and the extent of the tilting angles. Although some of these parameters were kept consistent (i.e., transmit pulse cycles and tilting angles), others, such as the lateral aperture, were not investigated. It is worth noting that the MS-550S has a larger lateral aperture (14.08mm) than either the L22-14vX (12.8 mm) or the L35-16vX (8.9 mm), which likely impacted the ULM reconstruction fidelity beyond the change in frequency. Likewise, the elevational beamwidth of each transducer is different, which alters the sampling volume and thus efficiency of MB localization.

ULM breaks the diffraction limit via localization of MBs, side-stepping the classic ultrasound trade-off between imaging resolution and penetration depth. Instead, the trade-off is between spatial resolution and data acquisition time. In this report, we found that increasing ultrasound frequency can still play a part in this milieu by enabling localization of higher MB concentrations, thus improving vessel saturation rate and confidence in microvascular reconstruction for neuroimaging research. This process is straightforward and accessible in the pre-clinical imaging space due to the prevalence of commercial high-frequency imaging probes.

## 5. Ending sections

### 5.1. Data and Code Availability

The data supporting the findings in the publication are available upon reasonable request.

### 5.2. Author Contributions

MRL and PS designed research; MRL, YW, BZL, ZH, DY, and YS performed research; MRL analyzed data; MRL and PS wrote the paper.

### 5.3. Funding

This study was partially supported by the National Institute of Biomedical Imaging and Bioengineering and the National Institute of Neurological Disorders and Stroke of the National Institutes of Health under grant numbers R21EB030072 and R56NS131516, by the National Science Foundation CAREER Award 2237166, and by the Chan Zuckerberg Initiative Ben Barres Early Career Award. The content is solely the responsibility of the authors and does not necessarily represent the official views of the NIH and NSF. MRL was supported by a Beckman Institute Postdoctoral Fellowship.

## References

Bar-Zion, A., Solomon, O., Tremblay-Darveau, C., Adam, D., & Eldar, Y. C. (2018). SUSHI: Sparsity-Based Ultrasound Super-Resolution Hemodynamic Imaging. *IEEE Transactions on Ultrasonics*, Ferroelectrics, and Frequency Control, 65(12), 2365–2380. 10.1109/TUFFC.2018.2873380

Belgharbi, H., Porée, J., Damseh, R., Perrot, V., Milecki, L., Delafontaine-Martel, P., Lesage, F., & Provost, J. (2023). An Anatomically Realistic Simulation Framework for 3D Ultrasound Localization Microscopy. *IEEE Open Journal of Ultrasonics*, Ferroelectrics, and Frequency Control, 3, 1–13. IEEE Open Journal of Ultrasonics, Ferroelectrics, and Frequency Control. 10.1109/OJUFFC.2023.3235766

Bourquin, C., Poree, J., Lesage, F., & Provost, J. (2022). In Vivo Pulsatility Measurement of Cerebral Microcirculation in Rodents Using Dynamic Ultrasound Localization Microscopy. IEEE Transactions on Medical Imaging, 41(4), 782–792. 10.1109/TMI.2021.3123912

Chabouh, G., Denis, L., Bodard, S., Lager, F., Renault, G., Chavignon, A., & Couture, O. (2024). Whole organ volumetric sensing Ultrasound Localization Microscopy for characterization of kidney structure. *IEEE Transactions on Medical Imaging*, PP. 10.1109/TMI.2024.3411669

Chang, H.-C., & Wang, L.-C. (2010). A Simple Proof of Thue’s Theorem on Circle Packing (arXiv:1009.4322). arXiv. 10.48550/arXiv.1009.4322

Chavignon, A., Hingot, V., Orset, C., Vivien, D., & Couture, O. (2022). 3D transcranial ultrasound localization microscopy for discrimination between ischemic and hemorrhagic stroke in early phase. Scientific Reports, 12(1), 14607. 10.1038/s41598-022-18025-x

Chen, X., Lowerison, M. R., Dong, Z., Han, A., & Song, P. (2022). Deep Learning-Based Microbubble Localization for Ultrasound Localization Microscopy. IEEE Transactions on Ultrasonics, Ferroelectrics, and Frequency Control, 69(4), 1312– 1325. 10.1109/TUFFC.2022.3152225

Christensen-Jeffries, K., Brown, J., Harput, S., Zhang, G., Zhu, J., Tang, M., Dunsby, C., & Eckersley, R. J. (2019). Poisson Statistical Model of Ultrasound Super-Resolution Imaging Acquisition Time. IEEE Transactions on Ultrasonics, Ferroelectrics, and Frequency Control, 66(7), 1246–1254. 10.1109/TUFFC.2019.2916603

Christensen-Jeffries, K., Browning, R. J., Tang, M.-X., Dunsby, C., & Eckersley, R. J. (2015). In vivo acoustic super-resolution and super-resolved velocity mapping using microbubbles. IEEE Transactions on Medical Imaging, 34(2), 433–440. 10.1109/TMI.2014.2359650

Couture, O., Besson, B., Montaldo, G., Fink, M., & Tanter, M. (2011). Microbubble ultrasound super-localization imaging (MUSLI). 2011 IEEE International Ultrasonics Symposium, 1285–1287. 10.1109/ULTSYM.2011.6293576

Couture, O., Hingot, V., Heiles, B., Muleki-Seya, P., & Tanter, M. (2018). Ultrasound Localization Microscopy and Super-Resolution: A State of the Art. IEEE Transactions on Ultrasonics, Ferroelectrics, and Frequency Control, 65(8), 1304– 1320. 10.1109/TUFFC.2018.2850811

Demené, C., Robin, J., Dizeux, A., Heiles, B., Pernot, M., Tanter, M., & Perren, F. (2021). Transcranial ultrafast ultrasound localization microscopy of brain vasculature in patients. Nature Biomedical Engineering, 5(3), 219–228. 10.1038/s41551-021-00697-x

Dencks, S., Piepenbrock, M., Schmitz, G., Opacic, T., & Kiessling, F. (2017). Determination of adequate measurement times for super-resolution characterization of tumor vascularization. 2017 IEEE International Ultrasonics Symposium (IUS), 1–4. 10.1109/ULTSYM.2017.8092351

Desailly, Y., Couture, O., Fink, M., & Tanter, M. (2013). Sono-activated ultrasound localization microscopy. Applied Physics Letters, 103(17), 174107. 10.1063/1.4826597

Desailly, Y., Pierre, J., Couture, O., & Tanter, M. (2015). Resolution limits of ultrafast ultrasound localization microscopy. Physics in Medicine and Biology, 60(22), 8723–8740. 10.1088/0031-9155/60/22/8723

Desailly, Y., Tissier, A.-M., Correas, J.-M., Wintzenrieth, F., Tanter, M., & Couture, O. (2017). Contrast enhanced ultrasound by real-time spatiotemporal filtering of ultrafast images. Physics in Medicine and Biology, 62(1), 31–42. 10.1088/1361-6560/62/1/31

Errico, C., Pierre, J., Pezet, S., Desailly, Y., Lenkei, Z., Couture, O., & Tanter, M. (2015). Ultrafast ultrasound localization microscopy for deep super-resolution vascular imaging. Nature, 527(7579), 499–502. 10.1038/nature16066

Heiles, B., Chavignon, A., Bergel, A., Hingot, V., Serroune, H., Maresca, D., Pezet, S., Pernot, M., Tanter, M., & Couture, O. (2022). Volumetric Ultrasound Localization Microscopy of the Whole Rat Brain Microvasculature. *IEEE Open Journal of Ultrasonics*, Ferroelectrics, and Frequency Control, 2, 261–282. 10.1109/OJUFFC.2022.3214185

Hingot, V., Chavignon, A., Heiles, B., & Couture, O. (2021). Measuring Image Resolution in Ultrasound Localization Microscopy. IEEE Transactions on Medical Imaging, 40(12), 3812–3819. 10.1109/TMI.2021.3097150

Hingot, V., Errico, C., Heiles, B., Rahal, L., Tanter, M., & Couture, O. (2019). Microvascular flow dictates the compromise between spatial resolution and acquisition time in Ultrasound Localization Microscopy. Scientific Reports, 9(1), 2456. 10.1038/s41598-018-38349-x

Huang, C., Lowerison, M. R., Trzasko, J. D., Manduca, A., Bresler, Y., Tang, S., Gong, P., Lok, U.-W., Song, P., & Chen, S. (2020). Short Acquisition Time Super-Resolution Ultrasound Microvessel Imaging via Microbubble Separation. Scientific Reports, 10(1), Article 1. 10.1038/s41598-020-62898-9

Jaqaman, K., Loerke, D., Mettlen, M., Kuwata, H., Grinstein, S., Schmid, S. L., & Danuser, G. (2008). Robust single-particle tracking in live-cell time-lapse sequences. Nature Methods, 5(8), Article 8. 10.1038/nmeth.1237

Kim, J., Wang, Q., Zhang, S., & Yoon, S. (2021). Compressed Sensing-Based Super-Resolution Ultrasound Imaging for Faster Acquisition and High Quality Images. IEEE Transactions on Bio-Medical Engineering, 68(11), 3317–3326. 10.1109/TBME.2021.3070487

Kuo, I. Y., & Shung, K. K. (1994). High frequency ultrasonic backscatter from erythrocyte suspension. IEEE Transactions on Bio-Medical Engineering, 41(1), 29–34. 10.1109/10.277268

Lee, S. A., Leconte, A., Wu, A., Kinugasa, J., Poree, J., Linninger, A., & Provost, J. (2024). Functional Assessment of Cerebral Capillaries using Single Capillary Reporters in Ultrasound Localization Microscopy (arXiv:2407.07857). arXiv. 10.48550/arXiv.2407.07857

Lin, H., Wang, Z., Liao, Y., Yu, Z., Xu, H., Qin, T., Tang, J., Yang, X., Chen, S., Zhang, X., Chen, X., & Shen, Y. (2024). Lifespan Super-Resolution Ultrasound Imaging Reveals Temporal Evolution of Cerebrovascular Alterations in the 5×FAD Mouse Model of Alzheimer’s Disease: Correlation with Pathological Impairments (SSRN Scholarly Paper 4794470). 10.2139/ssrn.4794470

Lowerison, M. R., Huang, C., Kim, Y., Lucien, F., Chen, S., & Song, P. (2020). In Vivo Confocal Imaging of Fluorescently Labeled Microbubbles: Implications for Ultrasound Localization Microscopy. *IEEE Transactions on Ultrasonics*, Ferroelectrics, and Frequency Control, 67(9), 1811–1819. IEEE Transactions on Ultrasonics, Ferroelectrics, and Frequency Control. 10.1109/TUFFC.2020.2988159

Lowerison, M. R., Sekaran, N. V. C., Dong, Z., Chen, X., You, Q., Llano, D. A., & Song, P. (2024). Super-Resolution Ultrasound Reveals Cerebrovascular Impairment in a Mouse Model of Alzheimer’s Disease. Journal of Neuroscience, 44(9). 10.1523/JNEUROSCI.1251-23.2024

Lowerison, M. R., Sekaran, N. V. C., Zhang, W., Dong, Z., Chen, X., Llano, D. A., & Song, P. (2022). Aging-related cerebral microvascular changes visualized using ultrasound localization microscopy in the living mouse. Scientific Reports, 12(1), Article 1. 10.1038/s41598-021-04712-8

Renaudin, N., Demené, C., Dizeux, A., Ialy-Radio, N., Pezet, S., & Tanter, M. (2022). Functional ultrasound localization microscopy reveals brain-wide neurovascular activity on a microscopic scale. Nature Methods, 19(8), Article 8. 10.1038/s41592-022-01549-5

Renaudin, N., Pezet, S., Ialy-Radio, N., Demene, C., & Tanter, M. (2023). Backscattering amplitude in ultrasound localization microscopy. Scientific Reports, 13(1), 11477. 10.1038/s41598-023-38531-w

Shelton, S. E., Lee, Y. Z., Lee, M., Cherin, E., Foster, F. S., Aylward, S. R., & Dayton, P. A. (2015). Quantification of Microvascular Tortuosity during Tumor Evolution Using Acoustic Angiography. Ultrasound in Medicine & Biology, 41(7), 1896– 1904. 10.1016/j.ultrasmedbio.2015.02.012

Shin, Y., Lowerison, M. R., Wang, Y., Chen, X., You, Q., Dong, Z., Anastasio, M. A., & Song, P. (2024). Context-aware deep learning enables high-efficacy localization of high concentration microbubbles for super-resolution ultrasound localization microscopy. Nature Communications, 15(1), 2932. 10.1038/s41467-024-47154-2

Siepmann, M., Schmitz, G., Bzyl, J., Palmowski, M., & Kiessling, F. (2011). Imaging tumor vascularity by tracing single microbubbles. 2011 IEEE International Ultrasonics Symposium, 1906–1909. 10.1109/ULTSYM.2011.0476

Song, P., Manduca, A., Trzasko, J. D., & Chen, S. (2017a). Noise Equalization for Ultrafast Plane Wave Microvessel Imaging. IEEE Transactions on Ultrasonics, Ferroelectrics, and Frequency Control, 64(11), 1776–1781. 10.1109/TUFFC.2017.2748387

Song, P., Manduca, A., Trzasko, J. D., & Chen, S. (2017b). Ultrasound Small Vessel Imaging With Block-Wise Adaptive Local Clutter Filtering. IEEE Transactions on Medical Imaging, 36(1), 251–262. 10.1109/TMI.2016.2605819

Song, P., Rubin, J. M., & Lowerison, M. R. (2023). Super-resolution ultrasound microvascular imaging: Is it ready for clinical use? Zeitschrift Für Medizinische Physik, 33(3), 309–323. 10.1016/j.zemedi.2023.04.001

van Heel, M., & Schatz, M. (2005). Fourier shell correlation threshold criteria. Journal of Structural Biology, 151(3), 250–262. 10.1016/j.jsb.2005.05.009

van Sloun, R. J. G., Solomon, O., Bruce, M., Khaing, Z. Z., Wijkstra, H., Eldar, Y. C., & Mischi, M. (2021). Super-Resolution Ultrasound Localization Microscopy Through Deep Learning. IEEE Transactions on Medical Imaging, 40(3), 829–839. 10.1109/TMI.2020.3037790

Yan, J., Zhang, T., Broughton-Venner, J., Huang, P., & Tang, M.-X. (2022). Super-Resolution Ultrasound Through Sparsity-Based Deconvolution and Multi-Feature Tracking. IEEE Transactions on Medical Imaging, 41(8), 1938–1947. 10.1109/TMI.2022.3152396

Zhang, Z., Hwang, M., Kilbaugh, T. J., Sridharan, A., & Katz, J. (2022). Cerebral microcirculation mapped by echo particle tracking velocimetry quantifies the intracranial pressure and detects ischemia. Nature Communications, 13(1), 666. 10.1038/s41467-022-28298-5

